# Genome sequence of the ornamental plant *Aquilegia vulgaris* reveals the flavonoid biosynthesis gene repertoire

**DOI:** 10.1101/2024.12.16.628782

**Authors:** Julie Anne V. S. de Oliveira, Ronja Friedhoff, Katharina Wolff, Boas Pucker

## Abstract

*Aquilegia vulgaris* is a widespread ornamental plant. The species is well known for its extensive variation of different flower colors and can even be considered as a model species for the evolution of flower morphology. Anthocyanins are a major pigment group responsible for pigmentation of flowers in *A. vulgaris* and many plant species. Here, we report a highly continuous genome sequence of an European *Aquilegia vulgaris* plant displaying purple flowers and a genome sequence of a plant displaying white flowers. The corresponding annotation facilitates research on flower color and morphology evolution. Long-read alignments revealed different structural variants in an anthocyanidin synthase (*ANS*) gene, crucial for pigment biosynthesis, that could explain the white flower phenotype. Candidate genes for all steps in the core anthocyanidin biosynthesis were identified. The identification of a flavonoid 3’,5’ hydroxylase (*F3’5’H*), a gene essential for the biosynthesis of bluish delphinidin derivatives, corroborates previous reports about metabolites and transcripts in *A. vulgaris*.

## Introduction

*Aquilegia vulgaris*, also known as Columbine or Granny’s Bonnet, is cultivated as an ornamental plant in many gardens. This plant species is perennial and considered easy to grow. *A. vulgaris* is well known for a huge variation of different flower colors and can even be considered as a model species for the evolution of flower morphology (Kramer & Hodges, 2010). Flower color is often caused by anthocyanins, which are produced by one branch of the flavonoid biosynthesis. Carotenoids, which are independently produced, could also contribute to the flower color phenotype. PCR-based searches with degenerated primers and RACE have resulted in the identification of 34 genes associated with the flavonoid biosynthesis (Hodges & Derieg, 2009). Many of these genes appear to be single copy, thus providing a potential explanation for the phenotypic diversity that could result from loss-of-function mutations (Hodges & Derieg, 2009). Multiple cases of independent floral pigmentation loss have been studied in *Aquilegia* discovering low expression of the central anthocyanin biosynthesis genes dihydroflavonol 4-reductase (*DFR*) and anthocyanidin synthase (*ANS*) (Whittall *et al*., 2006). Anthocyanins have a wide range of functions in plants, including scavenging reactive oxygen species, protecting the plant against intense light, and attracting pollinators and seed dispersers (Grünig *et al*., 2025). In *Aquilegia*, anthocyanins have been investigated in the context of coadaptation between plants and pollinators (Taylor, 1984). Besides anthocyanins, numerous other floral traits have been investigated in the context of pollinator attraction (Edwards *et al*., 2021). A genome sequencing approach involving multiple *Aquilegia* plants from different geological regions and producing a genome sequence for *A. coerulea* revealed a high degree of allele sharing between different species (Filiault *et al*., 2018). Another study sequenced the genome of *A. oxysepala* var. *kansuensis* (Xie *et al*., 2020).

While previous studies were focused on lines from North America (Whittall *et al*., 2006; Hodges & Derieg, 2009), this study investigates a European *Aquilegia* plant. Seeds were collected from a purple flowering and a white flowering plant in a German garden to grow the individuals for the sequencing project.

## Results and Discussion

### Genome sequence and annotation of *Aquilegia vulgaris*

We report two genome sequences of the ornamental plant *Aquilegia vulgaris*. Based on ONT long read sequencing, two highly continuous assemblies were generated (**Table 1**). The assembly size of 324 Mbp for the purple variant and 314Mbp for the white variant is consistent with the results of a flow cytometry study, which reported a DNA amount of 0.39 pg for 1C (Pustahija *et al*., 2013). The N50 of 3 Mbp for the purple variant indicates a high continuity that is suitable for comparative genomics. The assembly completeness was indicated to be about 96% based on BUSCO genes. For the white flowering plant, the N50 of 19.5Mbp indicates a very high continuity and a completeness of about 97% based on detected BUSCO genes. The LTR Assembly Index (LAI) (Ou *et al*., 2018) was calculated for both assemblies to evaluate the continuity based on intact Long Terminal Repeat Retrotransposons (LTR-RTs). LAI scores less than 10 are considered ‘draft quality’ assembly, between 10 and 20 are ‘reference quality’, and above 20 is considered ‘gold quality’ (Ou *et al*., 2018). Both assemblies are considered reference quality, with LAI scores of 11 for the purple variant and 14 for the white variant, respectively.

**Table 1:** Statistics of the *Aquilegia vulgaris* genome sequence produced in this study and comparison against publicly available genome sequences of other *Aquilegia* species.

|  | *Aquilegia vulgaris*<br>(purple-this study) | *Aquilegia vulgaris*<br>(white-this study) | *A. coerulea*<br>'Goldsmith'<br>(Filiault et al., 2018) | *A. oxysepala*<br>var. *kansuensis*<br>(Xie et al., 2020) |
| --- | --- | --- | --- | --- |
| **Total assembly size** | 323.7 Mbp | 315.2 Mbp | 291.7 Mbp | 297.8 Mbp |
| **Number of sequences** | 324 (contigs) | 120 (contigs) | 2529 (scaffolds) | 852 (contigs) |
| **Maximal contig length** | 16.8 Mbp | 47.4 Mbp | - | 7.8 Mbp |
| **N50** | 3.1 Mbp | 37.1 Mbp | 3.1 Mbp | 2.2 Mbp |
| **BUSCO** | C:94.9%[S:81.7%,D:13.2%],F:1.8%,M:3.3%,n:2805,E:2.8% | C:96.1%[S:91.0%,D:5.1%],F:0.7%,M:3.1%,n:2805,E:31.3% | C:96.0%[S:91.3%,D:4.6%],F:2.2%,M:1.8%,n:2805,E:3.2% | - |
| **LAI** | 11.71 | 14.38 | - | - |

The structural annotation of all protein encoding genes facilitates studies of the genetic basis of flower color formation and other floral traits. Different approaches to identify protein encoding genes have been attempted (**Table 2**). GeMoMa generated the best structural annotation based on hints derived from various related plant species. A completeness check of the annotation with BUSCO revealed about 94% of all expected genes in the purple flowering plant and 95% in the white one, respectively. The total number of genes is comparable to the average number of genes in a plant species (Pucker & Brockington, 2018).

**Table 2:**
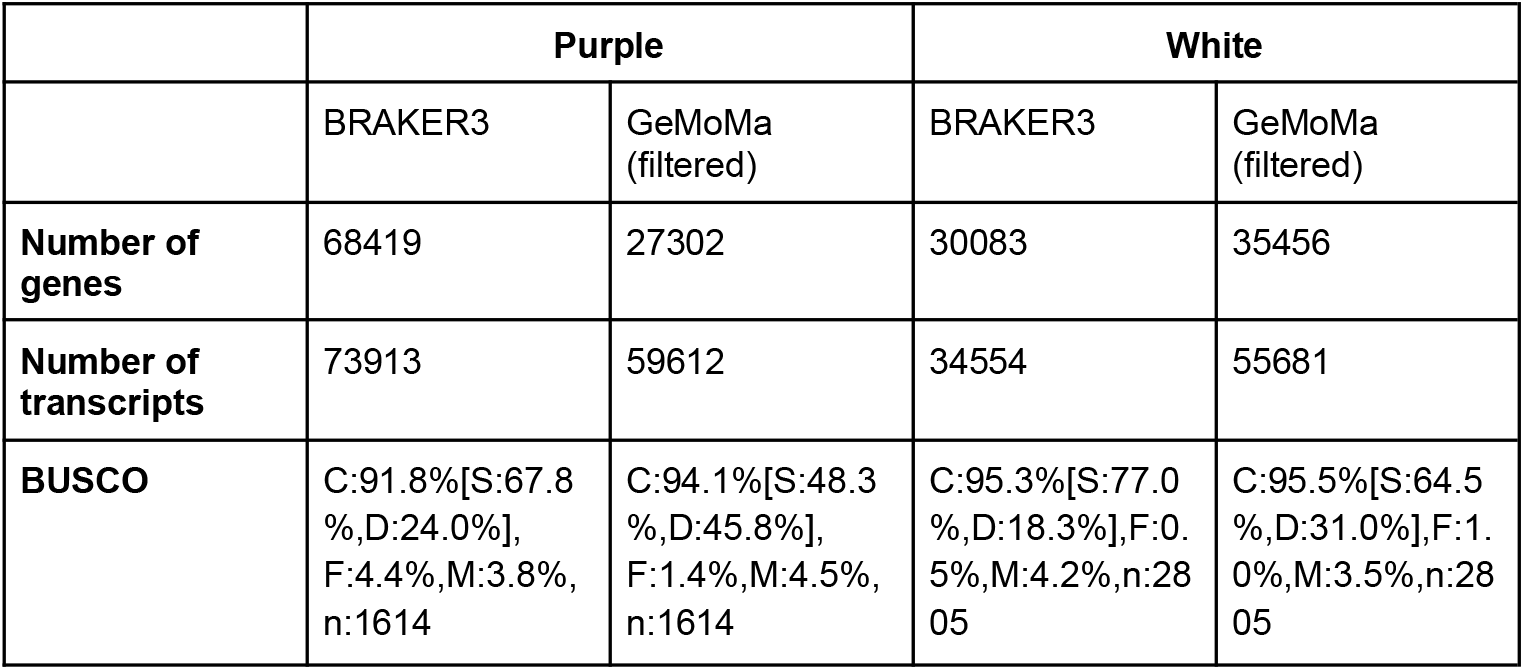
Comparison of different structural annotation approaches of *Aquilegia vulgaris*. BRAKER3 was run with RNA-seq hints. GeMoMa was supplied with RNA-seq hints and data sets of *Coptis chinensis* (GCA_015680905.1) (Liu *et al*., 2021), *Aquilegia coerulea* v3.1 (Acoerulea_322, GCA_002738505.1) (Filiault *et al*., 2018), *Thalictrum thalictroides* (GCA_013358455.1) (Arias *et al*., 2021), and *Ranunculus cassubicifolius (GCA_049309505*.*1)* (Karbstein *et al*., 2025). For the white *Aquilegia vulgaris*, GeMoMa was also supplied with datasets from the purple *Aquilegia vulgaris* generated in this study.

### Flavonoid biosynthesis in *Aquilegia vulgaris*

The flavonoid biosynthesis associated genes have been identified (**Fig. 2**, Additional file D). As expected for an anthocyanin-pigmented plant, genes for all steps in the anthocyanin biosynthesis were identified in the *A. vulgaris* gene set. This aligns with a previous study that reported 34 genes associated with the anthocyanin biosynthesis (Hodges & Derieg, 2009). The presence of F3’5’H is expected in this plant because delphinidin derivatives are believed to contribute to the purple coloration. In a previous study, *F3’5’H* of the closely related *A. buergeriana* was utilized in an attempt to engineer pigmentation of *Petunia hybrida* (Lee *et al*., 2023). The presence of F3’H and F3’5’H in *A. vulgaris* enables future investigations of the evolution of this cytochrome P450 gene family. It is already established that F3’5’H evolved multiple times independently from F3’H (Seitz *et al*., 2006, 2015), but the number of independent events remains unknown.

**Fig. 1.**
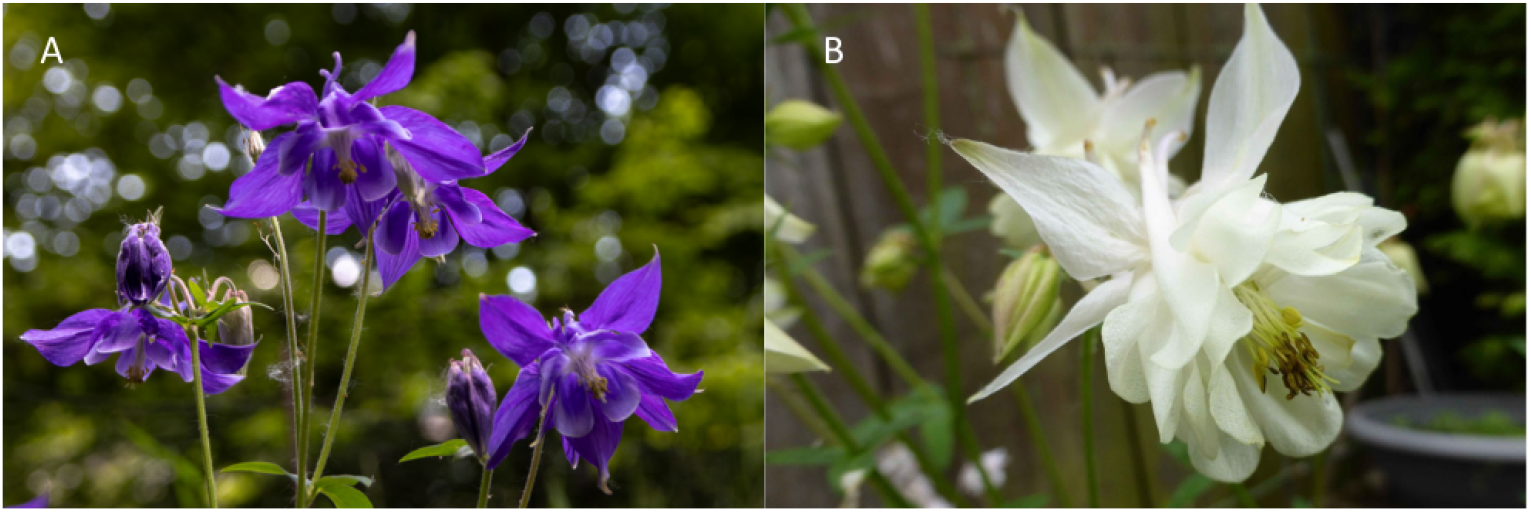
**A:** Purple flowering *Aquilegia vulgaris* plant. Photo credit: Jakob Horz. **B:** White flowering *Aquilegia vulgaris* plant. Photo credit: Pucker.

**Fig. 2:**
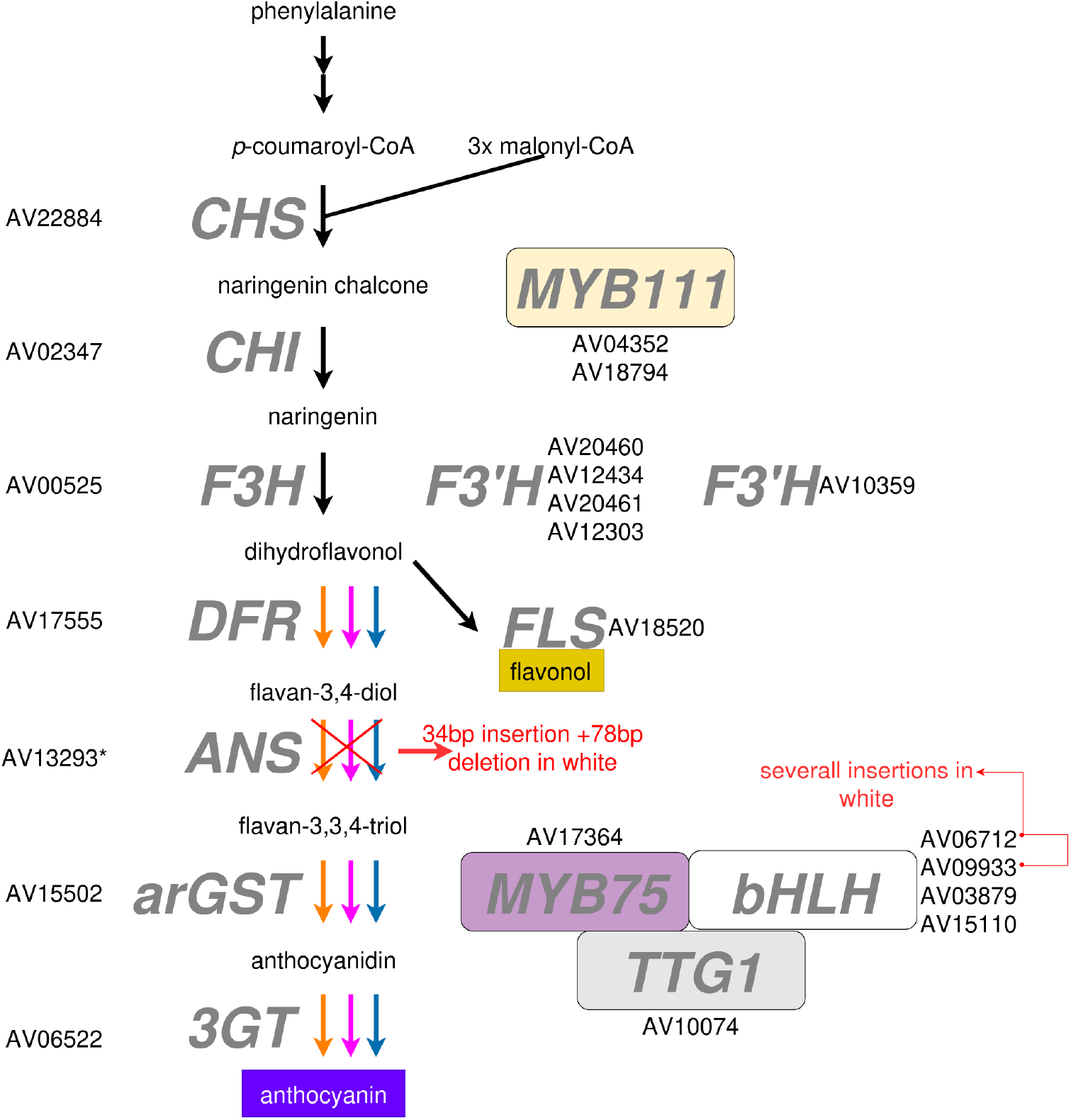
Candidate genes for steps in the flavonoid biosynthesis. The formation of different anthocyanins is indicated by arrows in the respective color: pelargonidin (orange), cyanidin (magenta), and delphinidin (blue).The variations observed only in the white flowering plant are marked with the red arrows. The layout is based on (Horz *et al*., 2025). *CHS*, chalcone synthase; *CHI*, chalcone isomerase; *F3H*, flavanone 3-hydroxylase; *F3’H*, flavonoid 3’-hydroxylase; F3’5’H, flavonoid 3’,5’-hydroxylase; *FLS*, flavonol synthase; *DFR*, dihydroflavonol 3-reductase; *ANS*, anthocyanidin synthase; *arGST*, anthocyanin-related glutathione S-transferase; *3GT*, UDP-dependent anthocyanidin 3-O-glucosyltransferase; *MYB*, Myeloblastosis; *bHLH*, basic helix-loop-helix; *TTG1, TRANSPARENT TESTA 1*.

An analysis of flavonoid biosynthesis gene expression was performed based on RNA-seq data of different seedling parts (Additional file 1, Additional file E).

The identified anthocyanin biosynthesis genes and corresponding transcription factors can serve as the basis for future studies exploring the genetic basis of flower color evolution. The huge diversity of flower colors in *Aquilegia* provides an excellent opportunity to conduct in-depth investigations of molecular mechanisms underlying flower color transitions (Kramer & Hodges, 2010). Besides wholesome loss events often caused by inactivation of transcription factors (Marin-Recinos & Pucker, 2024), more subtle changes can be explored. The contribution of flower colors to the adaptive radiation of plants can be investigated through comparison of the anthocyanin biosynthesis in different *Aquilegia* species. Interestingly, no candidate for an UDP-dependent anthocyanidin 3-O-glucosyltransferase was identified by KIPEs. A previous study reported the discovery of a flavonoid glucosyltransferase belonging to the UGT75B2 lineage (homolog of AT1G05530), but does not report an anthocyanin 3-O-glucosyltransferase of the UGT78D2 type (Tohge *et al*., 2005; Hodges & Derieg, 2009). While the absence of such a gene in the *A. vulgaris* gene set could be a technical artifact caused by the assembly or annotation process, it could also indicate a different glycosylation mechanism for anthocyanins. Several other glycosyltransferases were detected that might also accept anthocyanidins as substrate.

### Genetic variants responsible for pigmentation differences

Candidate genes in the anthocyanin biosynthesis pathway were examined manually through Integrative Genomics Viewer (IGV) (Robinson *et al*., 2023) with read alignments contrasting purple flowering and white flowering plants. A prominent feature in the *ANS* locus (AV13293), is a 34 bp insertion and a 78 bp deletion, both evident in the white flowering *A. vulgaris* (**Fig. 3**, Additional file A). These structural variants likely decrease ANS function, affecting anthocyanin production and yielding white flowers, as ANS catalyzes a key step in the pathway. The insertion and deletion appear homozygous in the white variant, potentially disrupting transcript splicing or coding sequence.

**Fig. 3:**
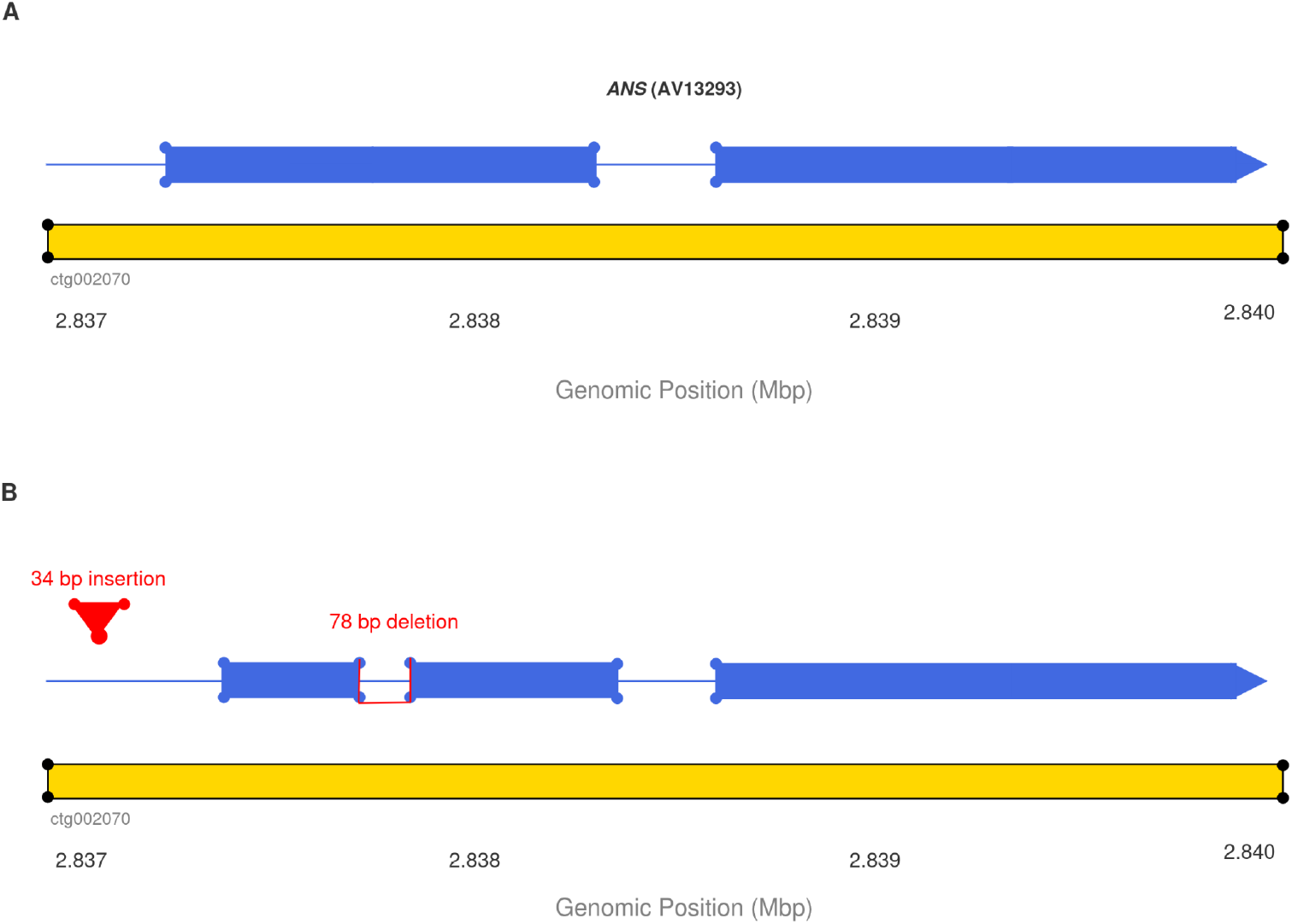
**A**. Reference *ANS* (AV13293) gene model in the purple genome sequence (ctg002070). Exons are shown as blue boxes, thin lines represent introns. **B**. *ANS* locus in the white-flowering genotype showing a 34 bp insertion (red triangle) at position 2,837,460 and a 78 bp deletion (red brackets) at position 2,838,082–2,838,159 within an exon. These white-specific variants are predicted to disrupt *ANS* function and anthocyanin biosynthesis.

Additionally, some bHLH transcription factors, that are key components of the flavonoid biosynthesis, carry specific variations only observed in the white *A. vulgaris*, this possibly contributes to a lack of pathway activation or could have been a secondary mutation events after loss of the *ANS* functionality. Notably, AV06712 (TT8-like bHLH) contains a 1,315 bp intron insertion that likely disrupts splicing (Additional file B), while AV09933 carries several small insertions (Additional file C). These variants might be able to compromise MYB-bHLH regulatory complexes thus lowering the activation of the anthocyanin biosynthetic genes. Previous studies have often attributed a lack of anthocyanin pigmentation to variants inactivating components of the MYB complex (Marin-Recinos & Pucker, 2024).

While validation in a larger population is required, these variants already present plausible explanations for the observed pigmentation differences. Especially the deletion in *ANS* associated with the loss of floral pigments aligns with known loss-of-function mutations affecting pigmentation in other plant species (Whittall *et al*., 2006; Rafique *et al*., 2016; Gonda *et al*., 2023; Horz *et al*., 2025). Meanwhile, genes in the flavonol (FLS) and proanthocyanidin (LAR, ANR) biosynthesis including their activator MYBs and TTG1 remain structurally intact, which suggests an anthocyanin specific biosynthesis block rather than a general flavonoid biosynthesis disruption. Loss of ANS activity might block the route towards ANR and epicatechins, but the path towards catechins via DFR and LAR appears unaffected. Future studies could explore the age of the white allele and its spread in the population. For example, a study in *Digitalis purpurea* revealed that the white allele predates human intervention thus suggesting that loss of flower coloration might not have a strong disadvantage in nature (Horz *et al*., 2025). Given the horticultural importance of *A. vulgaris*, it also appears plausible that this white allele emerged more recently and was propagated through breeding.

## Material and Methods

### DNA extraction and quality control

Plants were grown under long day conditions (16h light, 8 h darkness) in a plant cultivation room at approximately 20 °C. Prior to harvesting samples, the plants were incubated for one day in darkness to reduce starch content. Young leaves were harvested and homogenized by grinding in liquid nitrogen. High molecular DNA was extracted with a modified CTAB-based protocol as previously described (Siadjeu *et al*., 2020). Quality control of the DNA via NanoDrop measurement, agarose gel electrophoresis, and Qubit measurement was performed as previously described (Horz *et al*., 2025). Short DNA fragments were depleted with the Short Read Eliminator kit (Pacific Biosciences) following the supplier’s instructions.

### Nanopore sequencing

For the plant with purple flowers, libraries for the nanopore sequencing were prepared with the SQK-LSK109 ligation-based kit (Oxford Nanopore Technologies) using 1µg of DNA and following the suppliers’ instructions. Sequencing was conducted on a MinION-Mk1B with R9.4.1 flow cells. Upon blockage of a large proportion of nanopores, a wash step was performed followed by the loading of a fresh library to achieve optimal performance of the flow cell. Basecalling of the raw sequencing data was performed with Guppy v6.4.6 (ONT) on a GPU in the de.NBI cloud. Since the experiment took place at different times, for the white flowering plant, libraries for nanopore sequencing were prepared with the SQK-LSK114 ligation-based kit (Oxford Nanopore Technologies) using 1 µg of DNA and following the supplier’s instructions. Sequencing was performed on a MinION-Mk1B with R10.4.1 flow cells. The flow cell washing was performed in the same way. Basecalling of the raw sequencing data was performed with Dorado v1.0.2 (ONT) for the Purple variant and with Dorado v1.1.0 (ONT) for the White variant, both with high-accuracy model (dna_r10.4.1_e8.2_400bps_hac@v5.2.0) on a GPU in the de.NBI cloud.

### Genome sequence assembly

The genome sequence from the purple columbine was assembled with NextDenovo v2.5.2 (Hu *et al*., 2024) using the following settings genome_size = 370m and read_cutoff = 1k. And for the white, the genome sequence was assembled with Hifiasm-0.25.0-r726 (Cheng *et al*., 2021). Assembly statistics were calculated with a customized Python script (contig_stats3.py, (Meckoni *et al*., 2023). Contig names were cleaned to avoid technical issues in the following steps using the Python script clean_genomic_fasta.py (https://github.com/bpucker/GenomeAssembly). A screening process for contamination with bacterial or fungal sequences was conducted with assembly_wb_screen.py (https://github.com/bpucker/GenomeAssembly). BUSCO v5.8.2 (Manni *et al*., 2021) was run in the genome mode to assess the completeness of the assembly based on the eudicotyledons_odb12 reference data set. The LTR Assembly Index (LAI) (Ou *et al*., 2018) was calculated following a previously established workflow (de Oliveira *et al*., 2026).

### Genome sequence annotation

To generate hints for the structural annotation, RNA-seq data sets (Additional file 3) were retrieved from the Sequence Read Archive (Leinonen *et al*., 2011; Katz *et al*., 2022) with fastq-dump (NCBI, 2020). Read mappings of the purple variant were performed with STAR v2.7.4a (Dobin *et al*., 2013; Dobin & Gingeras, 2015). For the white flowering plant, HISAT2 v2.2.1 (Kim *et al*., 2019) was used for the RNA-seq read mappings. The BAM files containing the RNA-seq read mapping were passed to BRAKER3 (Gabriel *et al*., 2023) to enable the inference of hints for the gene prediction. GeMoMa v1.9 (Keilwagen *et al*., 2016, 2019) was run on both assembled genome sequences with the following data sets as hints: *Coptis chinensis* (GCA_015680905.1) (Liu *et al*., 2021), *Aquilegia coerulea* v3.1 (Acoerulea_322, GCA_002738505.1) (Filiault *et al*., 2018), *Thalictrum thalictroides* (GCA_013358455.1) (Arias *et al*., 2021), and *Ranunculus cassubicifolius* (Karbstein *et al*., 2025), for the white flowering plant, datasets from the purple *Aquilegia vulgaris* generated in this study was also used for GeMoMa. The initial annotation of the purple *A. vulgaris* was filtered with GeMoMa using the following criteria f=“start==‘M’ and stop==‘*’ and (isNaN(tie) or tie>0) and tpc>0 and aa>50” atf=“tie>0 and tpc>0”. An analysis of all predicted polypeptide sequences with BUSCO v3.0.2 (Simão *et al*., 2015) in protein mode was conducted to assess the annotation completeness. As for the white variant, the following criteria was used: f=“start==‘M’ and stop==‘*’ and aa>=30 and avgCov>0 and (isNaN(bestScore) or bestScore/aa>=2.5) and iAA>=0.5 and pAA>=0.5” atf=“iAA>0.9 and pAA>0.9 and sumWeight>1 and avgCov>100 and tpc==1 and tie==1”. To assess the completeness of the annotation of the predicted polypeptide sequences, BUSCO v5.8.2 was used in protein mode. Functional annotation terms were assigned to the predicted polypeptide sequences with the Python script construct_anno.py (Pucker & Iorizzo, 2023) based on the Araport11 annotation of *Arabidopsis thaliana* (Lamesch *et al*., 2012; Cheng *et al*., 2017). A detailed annotation of the flavonoid biosynthesis genes was performed with KIPEs v3 (Rempel *et al*., 2023) based on the flavonoid biosynthesis bait set v3.3.7. The anthocyanin biosynthesis regulating MYBs were identified with the MYB_annotator v1.0.1 (Pucker, 2022) and the anthocyanin biosynthesis regulating bHLHs were identified with the bHLH_annotator v1.0.4 (Thoben & Pucker, 2023). In the purple flowering plant, a TTG1 candidate was identified based on high sequence similarity in an initial search via BLASTp (Altschul *et al*., 1990, 1997) with TTG1 sequences from numerous plant species (Pucker *et al*., 2020). The candidate was validated through construction of a phylogenetic tree with FastTree v2.1.10 (Price *et al*., 2010) based on an alignment with MAFFT v7.453 (Katoh & Standley, 2013).

### Gene expression analysis for purple variant

All available RNA-seq data sets were processed with kallisto v0.44 (Bray *et al*., 2016) based on the coding sequences predicted for *A. vulgaris*. Customized Python scripts were deployed to merge all individual count tables into one final combined table (Pucker & Iorizzo, 2023). Violin plots visualizing the expression of flavonoid biosynthesis genes were generated with a Python script using the modules matplotlib, pandas, scipy, numpy, and seaborn (Hunter, 2007; Virtanen *et al*., 2020; Harris *et al*., 2020; Waskom, 2021; Pucker & Iorizzo, 2023; The pandas development team, 2024).

### Identification of sequence variants related to anthocyanin loss

Reads of the purple and white flowering plants (Additional file 2) were aligned against the reference genome sequence of the purple *A. vulgaris* with minimap2-2.29-r1283 (Li, 2021), using the default parameters for ONT long reads. The BAM file resulting from the mapping was indexed with SAMtools (Li *et al*., 2009). To analyze the candidate genes associated with the flavonoid biosynthesis, alignments were inspected withIntegrative Genomics Viewer (IGV) (Robinson *et al*., 2023).

## Data availability

All data sets underlying this study are publicly available. Sequencing data have been deposited at the European Nucleotide Archive (PRJEB63451, Additional file 2). The assembled genome sequence and corresponding annotation for the purple and white variant are available via bonndata (https://doi.org/10.60507/FK2/GKLMOG).

## Acknowledgements

This work was supported by the de.NBI Cloud within the German Network for Bioinformatics Infrastructure (de.NBI) and ELIXIR-DE (Forschungszentrum Jülich and W-de.NBI-001, W-de.NBI-004, W-de.NBI-008, W-de.NBI-010, W-de.NBI-013, W-de.NBI-014, W-de.NBI-016, W-de.NBI-022). We thank all members of the research group Plant Biotechnology and Bioinformatics for their discussion and support. We acknowledge support from Project DEAL and the University of Bonn for open access publication. We also thank all students participating in our ‘Data Literacy in Genome Research’ course. We thank Jakob Horz for providing a picture of our *Aquilegia vulgaris* plant.

## Author contributions

JAVSdO, KW and BP designed the experiment. JAVSdO and KW did the sequencing. JAVSdO, RF, and BP conducted the bioinformatic analyses. JAVSdO and BP wrote the manuscript. All authors reviewed the final version of the manuscript and consented to its submission.

## Additional Files

Additional file 1: Gene expression information inferred from RNA-seq data.

Additional file 2: Description of *Aquilegia vulgaris* nanopore sequencing data sets.

Additional file 3: Datasets used as hints for structural annotation.

Additional file A: IGV inspection of white flowering plant read mapping against purple flowering genome, showing the *ANS* locus (AV13293).

Additional file B: IGV inspection of the *TT8* locus (AV06712) in the mapping of reads from the white flowering plant against the genome sequence of the purple flowering plant.

Additional file C: IGV inspection of the *TT8* locus (AV09933) in the mapping of reads from the white flowering plant against the genome sequence of the purple flowering plant.

Additional file D: Flavonoid biosynthesis candidate genes in the annotation of the genome sequence belonging to the purple and white flowering plant, respectively.

Additional file E: Expression analysis of flavonoid biosynthesis candidate genes in the leaves of seedlings (a), the roots of seedlings (b), and adult leaves (c). Displayed expression values are log-transformed transcripts per million (TPMs).

