## Supplementary figures and images for "Genome sequence of the ornamental plant *Aquilegia vulgaris* reveals the flavonoid biosynthesis gene repertoire"

### Additional file A

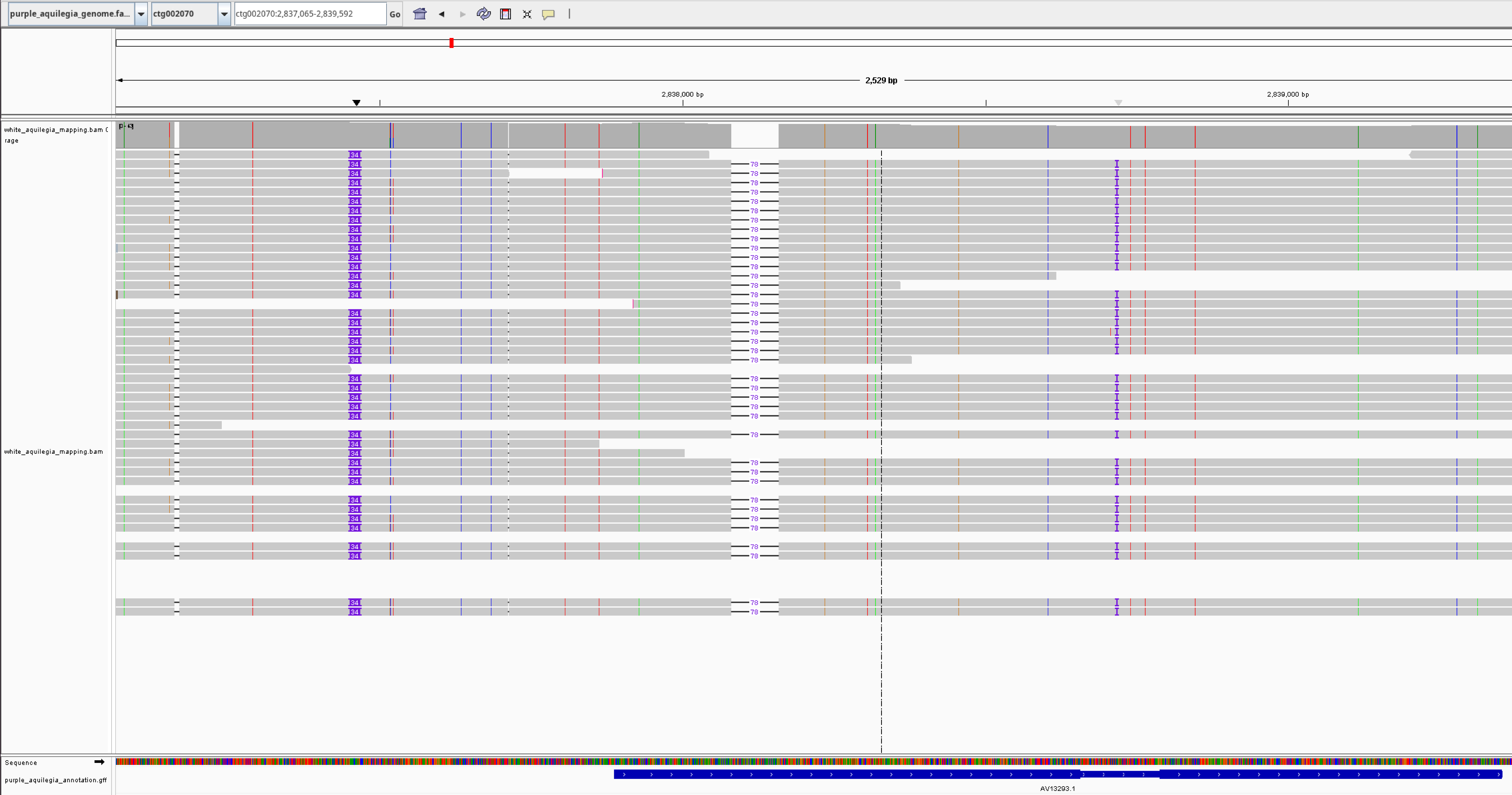

### Additional file B

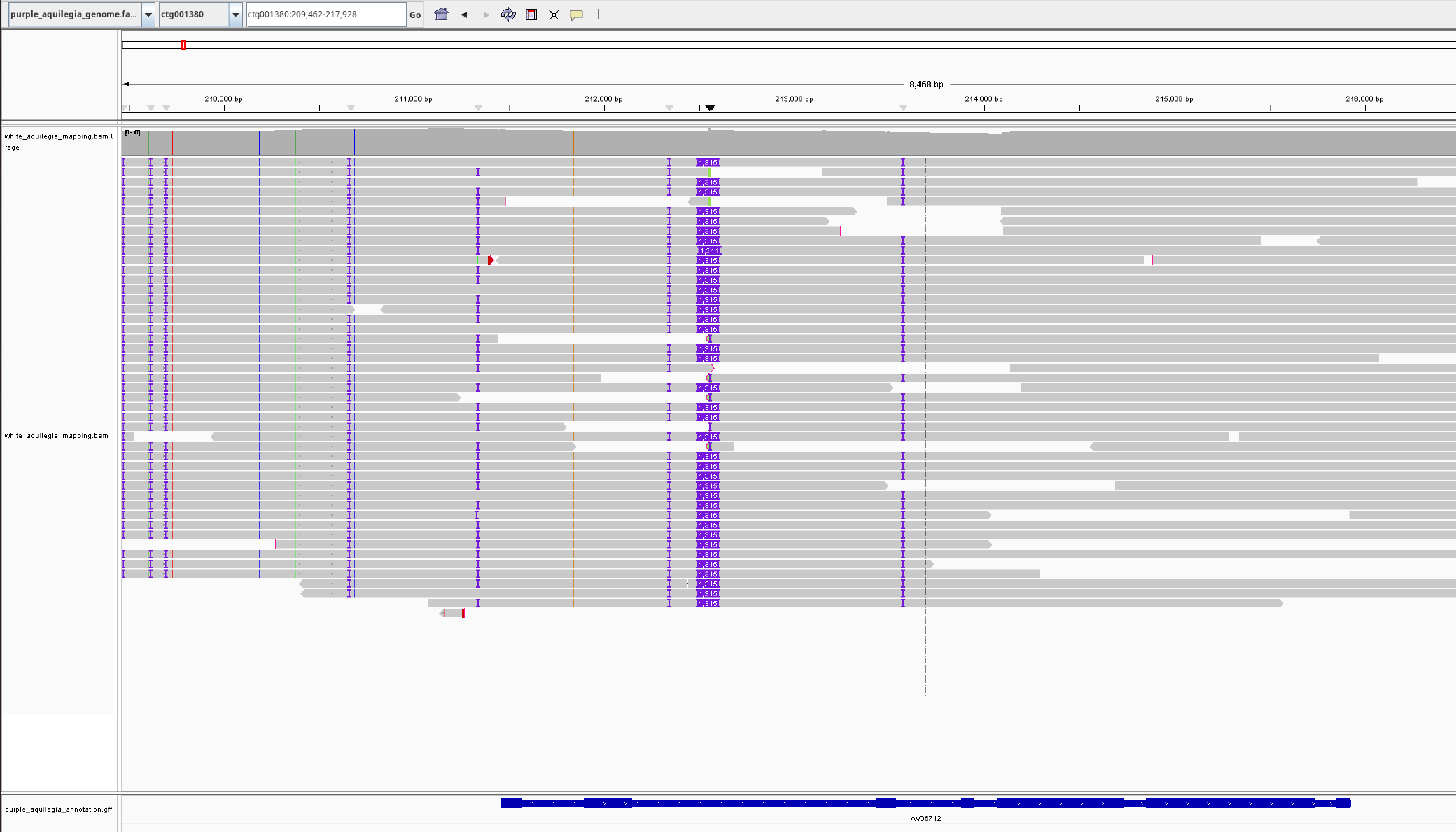

### Additional file C

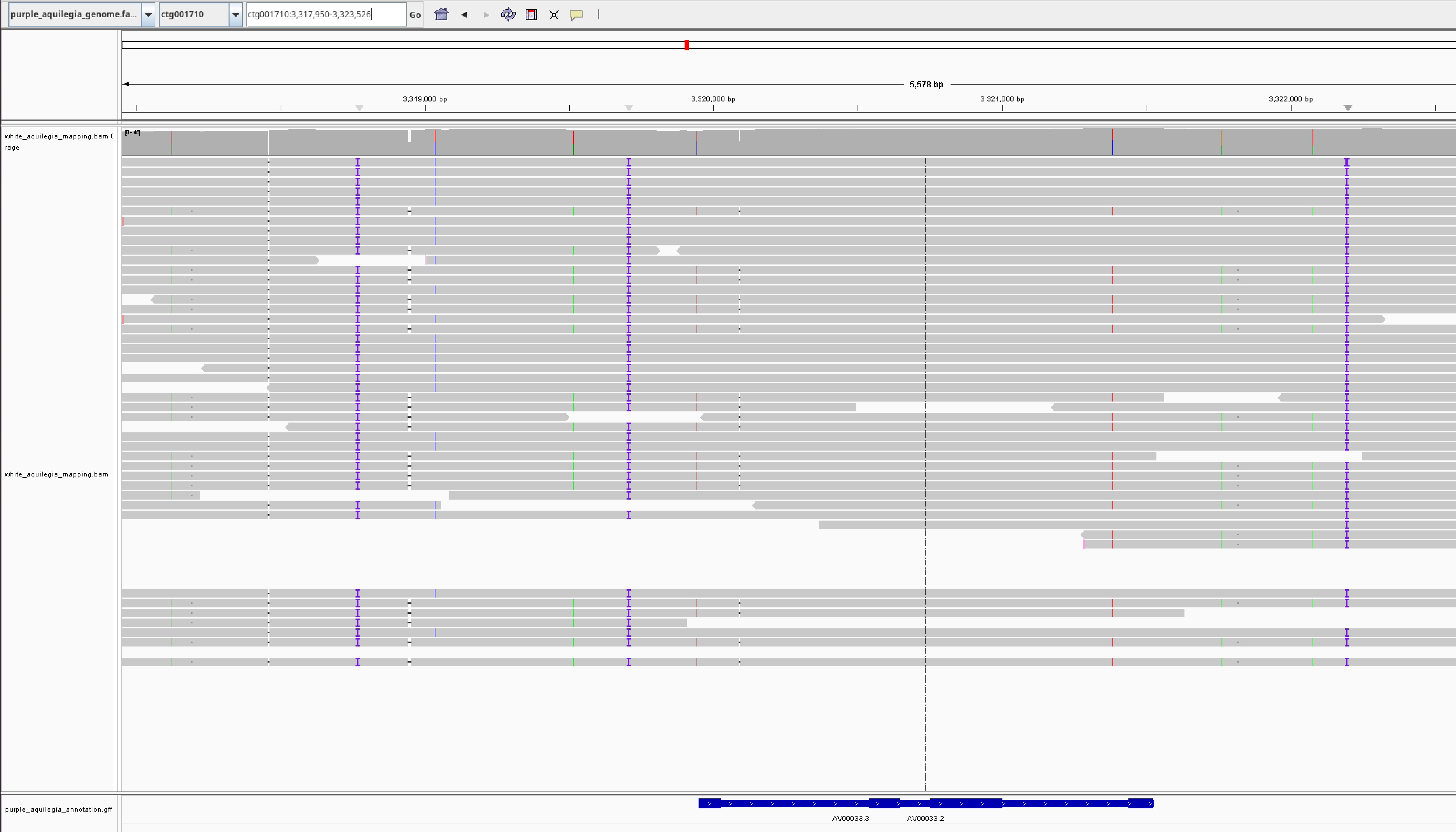

### Additional file E

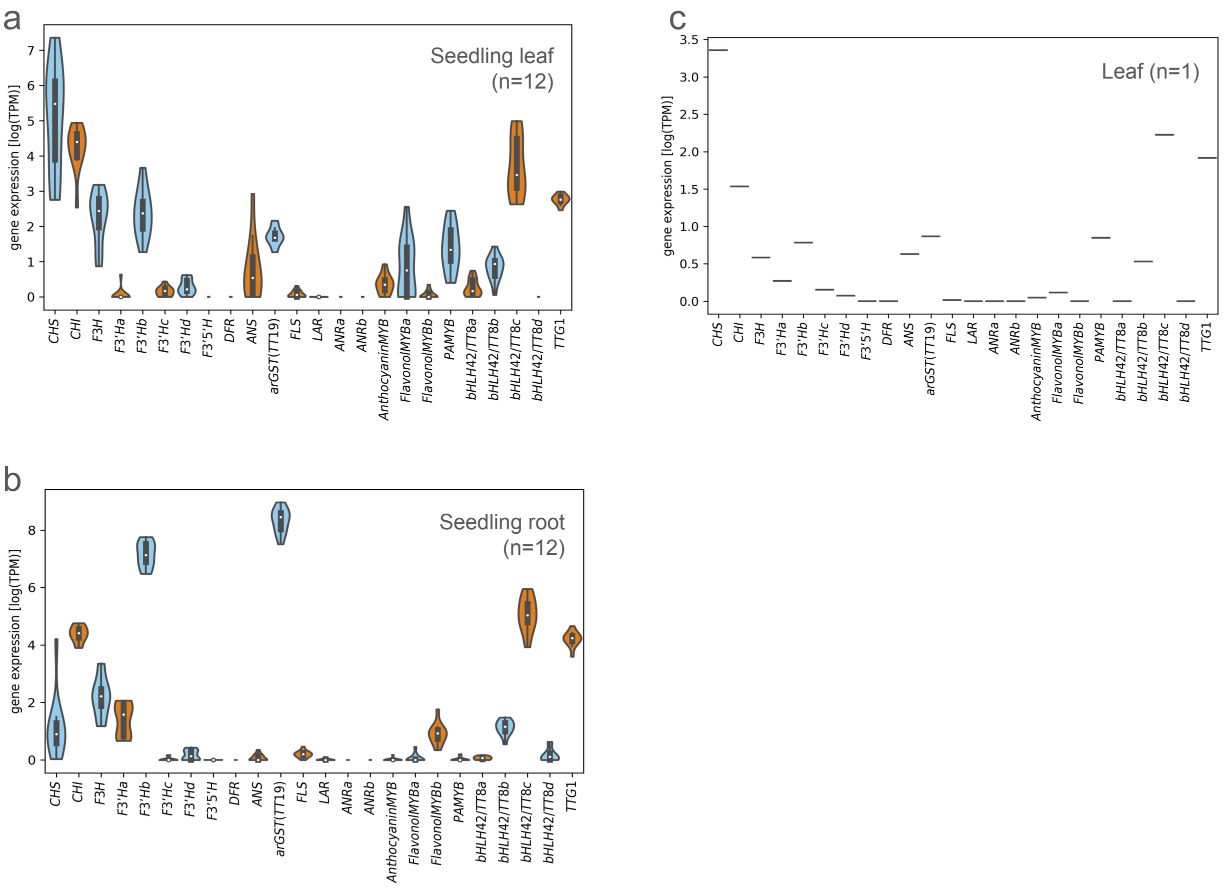
