## Additional file D for "Genome sequence of the ornamental plant *Aquilegia vulgaris* reveals the flavonoid biosynthesis gene repertoire"

| **Function** | **GeneID purple** | **GeneID white** |
|---|---|---|
| *CHS* | AV22884 | Avulgaris19207 |
| *CHI* | AV02347 | Avulgaris22446 |
| *F3H* | AV00525 | Avulgaris22543,Avulgaris09479 |
| *F3'H* | AV20460,AV12434,AV20461,AV12303 | Avulgaris19288 |
| *F3'5'H* | AV10359 | Avulgaris09087 |
| *DFR* | AV17555 | Avulgaris26447 |
| *ANS* | AV13293* | Avulgaris10383 |
| *arGST (TT19)* | AV15502 | Avulgaris19930 |
| *FLS* | AV18520 | Avulgaris25323 |
| *LAR* | AV11162 | Avulgaris06370 |
| *ANR* | AV09695, AV24619 | Avulgaris04343, Avulgaris04030 |
| Anthocyanin MYB | AV17364 | Avulgaris04259,Avulgaris26693,Avulgaris26702,Avulgaris26700,Avulgaris26701 |
| Flavonol MYB | AV04352,AV18794 | Avulgaris32261,Avulgaris23432 |
| Proanthocyanidin MYB | AV08268 | Avulgaris11039 |
| bHLH42/*TT8* | AV03879,AV06712,AV09933,AV15110 | Avulgaris03797,Avulgaris06613,Avulgaris14615,Avulgaris18441,Avulgaris24055,Avulgaris24641,Avulgaris33107,Avulgaris33460 |
| *TTG1* | AV10074 | Avulgaris16677 |
